# Female resource limitation does not make the opportunity for selection more female biased

**DOI:** 10.1101/748657

**Authors:** Ivain Martinossi-Allibert, Johanna Liljestrand Rönn, Elina Immonen

**Affiliations:** Department of Organismal Biology/Systematics Biology, Uppsala University, Norbyvägen 18D, SE-75236 Uppsala, Sweden; Department of Ecology and Genetics/Animal Ecology, Uppsala University, Norbyvägen 18D, SE-75236 Uppsala, Sweden; Department of Ecology and Genetics/Evolutionary Biology, Uppsala University, Norbyvägen 18D, SE-75236 Uppsala, Sweden

**Keywords:** Sex-specific selection, sexual selection, sexual dimorphism, sex-bias, opportunity for selection, laboratory settings

## Abstract

Environmental and physiological conditions affect how individual variation is expressed and translated into variance in fitness, the opportunity for natural selection. Competition for limiting resources can magnify variance in fitness and therefore selection, while abundance of resources should reduce it. But even in a common environment the strength of selection can be expected to differ across the sexes, as their fitness is often limited by different resources. Indeed most taxa show a greater opportunity for selection in males than in females, a bias often ascribed to intense competition among males for access to mating partners. This sex-bias could reverberate on many aspects of evolution, from speed of adaptation to genome evolution. It is unclear however, whether the sex-bias in opportunity for selection is robust to variations in environment or physiological condition that limit sex-specific resources. Here we test this in the model species *C. maculatus* by comparing female and male variance in relative fitness (opportunity for selection) under mate competition (i) with and without limitation of quality oviposition sites, and (ii) under delayed age at oviposition. Decreasing the abundance of the resource key to females or increasing their reproductive age was indeed challenging as shown by a reduction in mean fitness, however variance in fitness remained male-biased across the three treatments, with even an increased male-bias when females were limited by oviposition sites. This suggests that males remain the more variable sex independent of context, and that the opportunity for selection through males is indirectly affected by female-specific resource limitation.

## Introduction

Variation in fitness among individuals is what natural selection acts on. It can be partitioned into variation among individuals in their genetic makeup (breeding value), in their phenotypic condition subjected to environmental variation, and to the interaction between the two [1]. Therefore, the extent to which individual variation will be translated into variation in fitness visible to natural selection depends on context, through the availability of key developmental resources and the intensity of competition among individuals [2, 3]. For example, under abundant resources individual variation in resource acquisition should matter little to fitness, but if resources are scarce even slight differences in the acquisition traits may translate into large differences in fitness.

To see how variation in fitness translates into opportunity for selection, it is useful to think about a selection differential, which is the covariance between a trait and relative fitness [4, 1]. Variance in relative fitness then sets the upper limit for the strength of selection on any trait [5], as it represents the strength of selection on a trait that would covary perfectly with fitness. For this reason, variance in relative fitness has been called the opportunity for selection, often designated by ***I***. When a change in context affects the magnitude of fitness differences among individuals, it therefore affects variance in fitness and the opportunity for selection.

In sexually reproducing populations, males and females often have different reproductive strategies [6], which means that they can be limited by different resources. This results in a situation where a common environment can impose different challenges to the sexes, which should translate into sex-specific variance in fitness and sex-specific opportunity for selection. Indeed, sexual selection theory predicts that mating partners should often be a limiting resource for males, which together with natural selection should result in generally stronger net selection on males than on females [7]. If so, this could have far-reaching consequences in several aspects of evolutionary biology, such as speed of purging of deleterious mutations [8], speed of adaptation to novel environments [9], rate of evolution of sex-specific traits, or genome structure and evolution [10]. For example, the purging of deleterious mutations through selection on males, at the benefit of both sexes, has been proposed as one of the mechanism explaining the maintenance of sexual reproduction itself [8, 11]. Sex-biased fitness variance can also lead to different effective population sizes in the sexes, which can cause asymmetries in the genetic diversity of the sex chromosomes relative to autosomes [12, 13, 14].

Empirical works investigating patterns of sex-specific selection shows that in many species variance in reproductive success is indeed male-biased, e.g. [15, 16, 17, 18, 19], but not in all (reviewed in [20]). In their resent meta-analysis, Janicke et al. [21] gathered sex-specific estimates of variance in reproductive success and other selection metrics from 66 species in 72 studies, all from wild populations. Their work showed that, although there is variation across taxa and some species show female-biased selection or no sex-bias, the general trend is for male-biased selection (as measured with male-biased variance in reproductive success). In 2018, Singh and Punzalan [22] collated data from sex-specific estimates of phenotypic selection on traits (selection gradients), again in wild populations. With 865 estimates, they detected a general male-bias in selection, mostly driven by traits related to mating success. These two comprehensive studies therefore clearly support the hypothesis that there should be a general male-bias in selection, with some evidence indicating that this trend may be due to sexual selection specifically. However, these two studies have also revealed tremendous variability across taxa, and the source of this variability is still poorly understood.

If male fitness is expected to generally be more variable because of sexual selection, there are also many reasons for female fitness to exhibit high levels of variance. First, in some species they do experience strong sexual selection (reviewed in [23]), but there are also many other sources of fitness variation depending on the ecology of each species, such as competition for nutritional resources, nesting or oviposition sites [24]. The context in which selection is measured greatly matters, as the limitation of specific resources can magnify or shrink fitness differences among individuals. Because the sexes are sensitive to different limiting resources, variation in environmental conditions could unveil variation in fitness differently in the sexes, which has rarely been experimentally studied (but see [3]). Here we tested this hypothesis, and thus robustness of the pattern of male-biased opportunity for selection, by measuring sex-specific variance in relative fitness using three experimental conditions designed to specifically challenge female fitness. We predicted that conditions that limit female-specific resources should result in a more female-biased opportunity for selection. Understanding better female-specific environmental limitations should further our understanding of the natural variation in sex-specific patterns of selection.

To do this we used the seed beetle *Callosobruchus maculatus*, a widely used laboratory system for sexual selection studies [25]. We compared sex-specific variance in relative fitness (the opportunity for selection) in three different treatments: under a competitive context allowing sexual competition on both sexes and ad libitum oviposition substrate offered to females (control treatment, CT), under a heterogeneous-environment treatment (HT), presenting individuals with the context of sexual competition but with an oviposition substrate of heterogeneous quality; and an ageing treatment (AT) in which females were challenged physiologically to prolong their age at oviposition. This last treatment was chosen to challenge individuals through ageing, which is known to affect the sexes differently in *C. maculatus* [26, 27, 28]. For example, eggs from older mothers are less likely to hatch, while there are no detectable effects of paternal age on offspring phenotype [26]. Fertility also declines much more rapidly with age in females than in males [28]. Ageing therefore represents a greater challenge for female fitness in this species, which may result in more female-biased opportunity for selection. Moreover, both the HT and AT treatment should be relevant to the ecology of *C. maculatus* that, as a bean beetle, is dependent on patchy bean seeds as the only larval food resource, without which the females do not even lay eggs. Females have evolved a great capacity to detect a high quality bean resource as their oviposition site [29, 30]. The HT treatment thus provides a challenge that can reveal variation in this crucial ability for female fitness, while the AT treatment represents a situation faced by individuals required to postpone reproduction in the absence of available bean resources. We estimated the strength of selection as the opportunity for selection :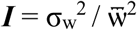, where σ_w_ is the standard deviation in fitness and 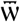 the mean fitness [5].

We find that mean fitness, measured as the number of adult offspring recruited to the next generation, was lower in both HT and AT treatments compared to the control, indicating they were generally challenging conditions. Interestingly, individual offspring produced by older parents (i.e. AT treatment) were heavier than ones from either of the other treatments, suggesting that this particular stressor induced a change in the offspring resource allocation strategy. Finally, the opportunity for selection was consistently higher in males than in females, and the male-bias was even stronger under oviposition site limitation (i.e. HT), suggesting that this sex-specific trend is not only robust to the context but that male variation can be indirectly affected through interaction with females.

## Methods

### Study organism and population

The seed beetle *Callosobruchus maculatus* is a facultative aphagous pest species found in grain storages and fields across West Africa and Asia. Its reproductive cycle, which typically spans over about a month, starts by adults laying eggs on the surface of beans (for example the black-eyed bean *Vigna ungulata* used in the present study), after which larvae burrow and develop inside the beans until they emerge as reproductively mature adults.

The study population originates from a natural population sampled in Lome, Togo (06°10#N 01°13#E) in 2010. It has been kept under laboratory conditions since then (29°C, 12:12 light cycle, 50% humidity) with a constant population size of approximately 400-500 individuals. Fitness assays were also performed under laboratory conditions (29°C, 12:12 light cycle, 50% humidity).

### Experimental design

#### Fitness assays

Fitness was measured in lifetime competitive assays were one focal individual was placed together with a competitor of the same sex and two mating partners of the opposite sex inside a 9cm petri dish. The environment inside the dish varied according to the treatment (see experimental treatments below). At the start of the experiment all individuals were adult virgins collected less than 24 hours after emergence from the beans. The competitor individual was sterilized by gamma radiation (100Gy), a commonly used method in the seed beetles that allows the competitor individual to compete for matings and achieve fertilizations, but insures that zygotes fertilized by the competitor will not develop due to the high number of double-stranded breaks in the embryo DNA caused by the irradiation [31, 32]. The four individuals were left to interact during their lifetime and offspring were counted as emerged adults of the next generation. A female fitness assay included one focal female, one sterilized female, and two male partners. The same design was used for the male fitness assays, which included one focal male, one sterilized male and two female partners.

#### Experimental treatments

Our study included three treatments, aimed to create different reproductive challenges for the sexes.

The control treatment (CT) represents the laboratory setting classically used in *C. maculatus* studies: a 9cm petri dish with ad libitum black-eyed bean (27g, approximately 130 beans). While male fitness variation can be manifested through pre- and post-mating sexual competition, for females this environment likely represents less challenges. Their oviposition substratum, the bean, is directly available, in a high and consistent quality, and in non-limiting quantity.

The heterogeneous environment treatment (HT) was designed to directly challenge females in their ability to discriminate quality oviposition sites. Each petri dish was filled with beans of variable quality: 15 high quality beans (3-4 grams) and the remainder of poor quality for a total of 27g as well. The low-quality beans were produced by letting a stock population of *C. maculatus* use the beans for larval development, resulting in bored beans that provide less resources for offspring to develop on.

The ageing treatment (AT) was designed to challenge females in their ability to withhold their reproduction until a suitable oviposition site is available. This treatment bears ecological relevance to a scenario where high-quality oviposition sites are exhausted upon female hatching, requiring prolonged periods of searching for suitable sites. In this treatment, the four individuals were first placed in an empty dish and left to interact for 48h, after which ad libitum (27g) high quality beans were added.

#### Sex-specific variance in fitness

To measure sex-specific variance in fitness we used competitive fitness assays including four individuals: the focal individual (female for a female assay and a male for a male assay), a sterile competitor of the same sex, and two potential mating partners of the opposite sex. Since all individuals originate from the same population, all the non-sterile individuals (one focal individual and two mating partners) will contribute to the final estimate of variance in fitness. For example, variance measured from female assays will be composed of a component due to the focal female present in each assay, but also of a component due to the two males present as potential mating partners. We considered the contribution of mating partners for estimating the sex-specific variance in fitness under the following premise. As the contribution of the mating partners is shared between two individuals, but the contribution of the focal individual relies solely on one in each assay, the focal sex contributes fully to the variance in fitness while the mating partners’ contribution is halved, so that:

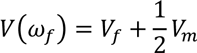

And,

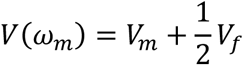

Where *V*(*ω*_*m*_) and *V*(*ω*_*f*_) are the variances estimated from female and male fitness assays respectively, and *V*_*f*_ and *V*_*m*_ are the female and male components of these variances. This premise stems from the assumption that the contributions of both parents to fitness are additive, and that breeding values of males and females are normally distributed.

If we call *F* the female breeding value and suppose that it follows a normal distribution with mean 1 and standard deviation 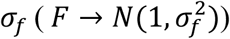, and call *M* the male breeding value and suppose 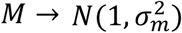, we can describe the fitness of a female assay as :

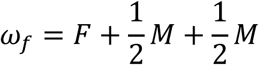

And of a male assay as:

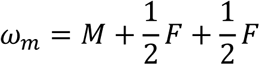

This is because the focal sex contributes fully (*F* or *M* in the female or male assay) while the mating partners share their reproductive output between the focal individual and the sterile competitor, giving on average one half each to the focal individual. The contributions of the mating partners are normally distributed with 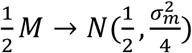 and 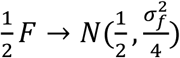. *ω*_*f*_ and *ω*_*f*_ are then also normally distributed, with variances:

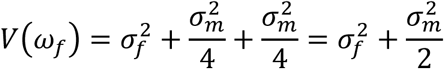

And

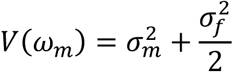

where we see that the male variance component is halved in the female assays and *vice versa* in the male assays. The ratio of *V*(*ω*_*m*_)/*V*(*ω*_*f*_) should therefore be a reliable indication of the sex bias in the opportunity for selection.

### Statistics

#### Mean fitness

The effect of experimental treatments on mean fitness (offspring number) was analyzed using a linear mixed-model, as implemented in the lme4 package (version 1.1-18-1, [33]) for R (version 3.5.1, [34]), taking into account normal distribution of the data. Experimental treatment, sex of the focal individual and their interaction were specified as fixed effect and date of the fitness assay as a random effect.

#### Individual offspring weight

The effect of experimental treatments on individual offspring weight was analyzed using a linear mixed-model, as implemented in the lme4 package for R, taking into account normal distribution of the data. Experimental treatment, sex of the focal individual and their interaction were specified as fixed effect and the date of the fitness assay as a random effect.

#### Sex-specific variance in fitness

A Bayesian model, as implemented in the MCMCglmm package (version 2.26, [35]) for R, was used to estimate components of variance in fitness attributed to each sex by experimental treatment combination. Because opportunity for selection is the variance in relative fitness, fitness data was mean standardized so that each sex by treatment subset had a mean of one prior to this analysis. The model was then specified with assay date as a random effect and the total phenotypic variance estimated for each sex by experimental treatment combination (*idh* structure not allowing for covariances to be estimated). For each experimental treatment, the log ratio of the posterior distributions for male and female variances were then computed, giving a mean log ratio and 95% confidence intervals.

## Results

### Mean fitness

Mean fitness (offspring number) differed among all experimental treatments (Table 1), being highest in the CT followed by the HT and finally the AT (Figure 1). The treatment differences from each other were confirmed by post-hoc tests (Tukey’s post-hoc, CT-AT: HSD=8.6, p<0.001, CT-HT: HSD=2.3, p=0.024, AT-HT: HSD=6.0, p<0.001, corrected for multiple testing with the Holm-Bonferroni method). A weak main effect of sex was also detected (Table1, Figure1) with males having slightly overall higher mean offspring number but there was no sex by treatment interaction. These result indicate that the HT and AT treatments were indeed challenging, with respectively 14% and 36% reduction in mean fitness compared to the control, and that the AT was more stressful than the HT.

**Table 1.** Anova table for a linear mixed model with offspring number as a response variable. Type III test. Date of the fitness assay was estimated as random effect.

| Effect | Chi square | df | p.value |
| --- | --- | --- | --- |
| Intercept | 275 | 1 | <0.001 |
| Treatment | 78.4 | 2 | <0.001 |
| Sex | 4.67 | 1 | 0.03 |
| Treatment by Sex | 3.53 | 2 | 0.17 |

**Figure 1.**
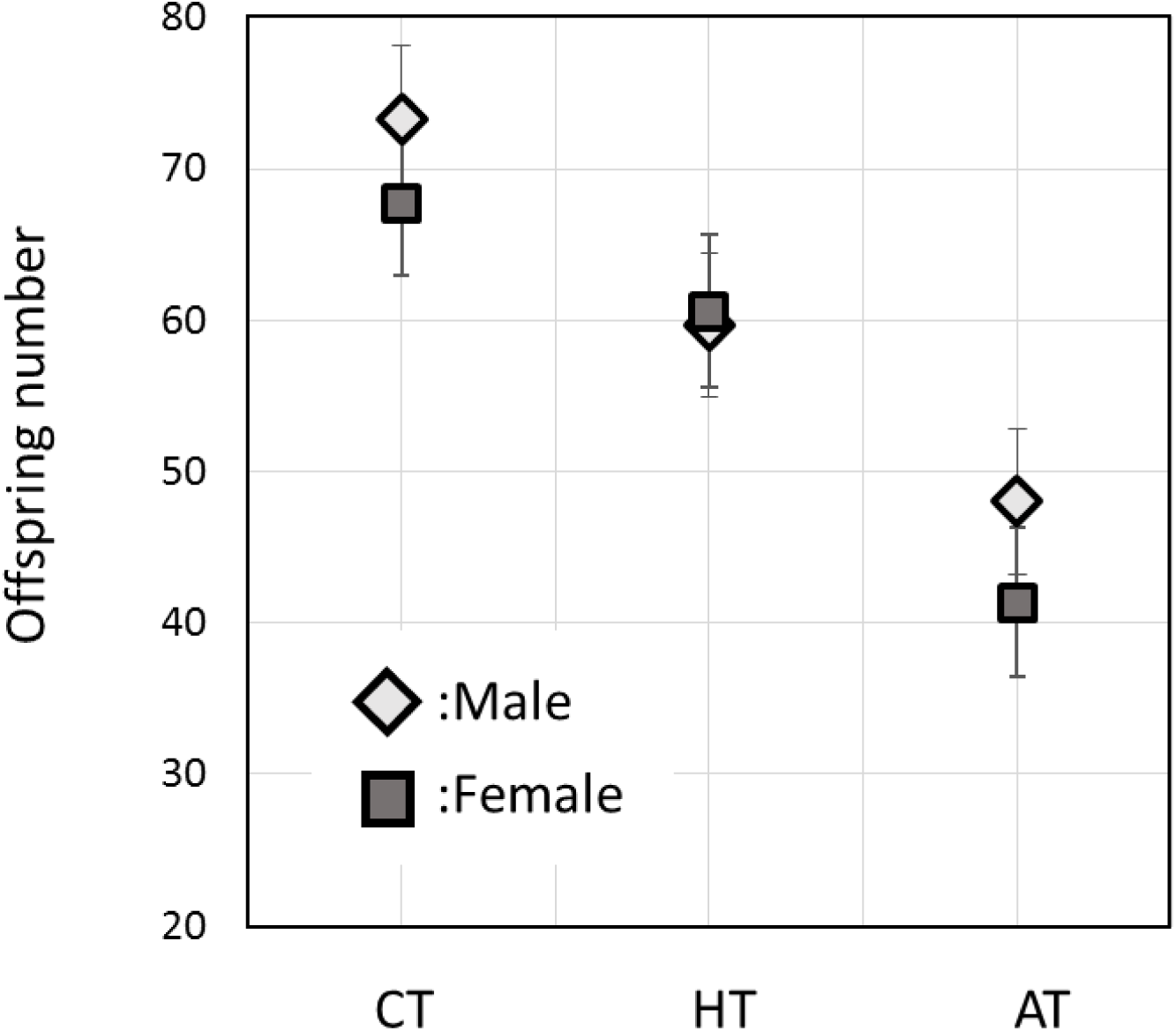
Mean fitness for each sex and experimental treatment. Mean fitness (adult offspring number) and 95% confidence limits (linear mixed model estimates) are given for each treatment: Control (CT), Heterogeneous treatment (HT) and Ageing treatment (AT). Female values are given by dark shaded squares and male values by light shaded diamonds.

### Total and Individual offspring weight

Mean total offspring weight differed among the experimental treatments (Table2): the CT had the highest total weight, while the HT and AT showed no difference (Figure 2a, Tukey’s post-hoc: CT-AT: HSD=3.5, p=0.001, CT-HT: HSD=2.9, p=0.009, AT-HT: HSD=0.55, p=0.58, corrected for multiple testing with the Holm-Bonferroni method). Thus, the HT and AT treatment had a different mean number of offspring, but the same mean total offspring weight. This is achieved by individuals from the AT treatment producing larger offspring (Figure 2b). More particularly, individual offspring weight was higher in the AT compared to both other treatments which did not differ from each other (Table3, Tukey’s post-hoc: CT-AT: HSD=7.6, p<0.001, CT-HT: HSD=1.8, p=0.07, AT-HT: HSD=9.2, p<0.001, corrected for multiple testing with the Holm-Bonferroni method).

**Figure 2.**
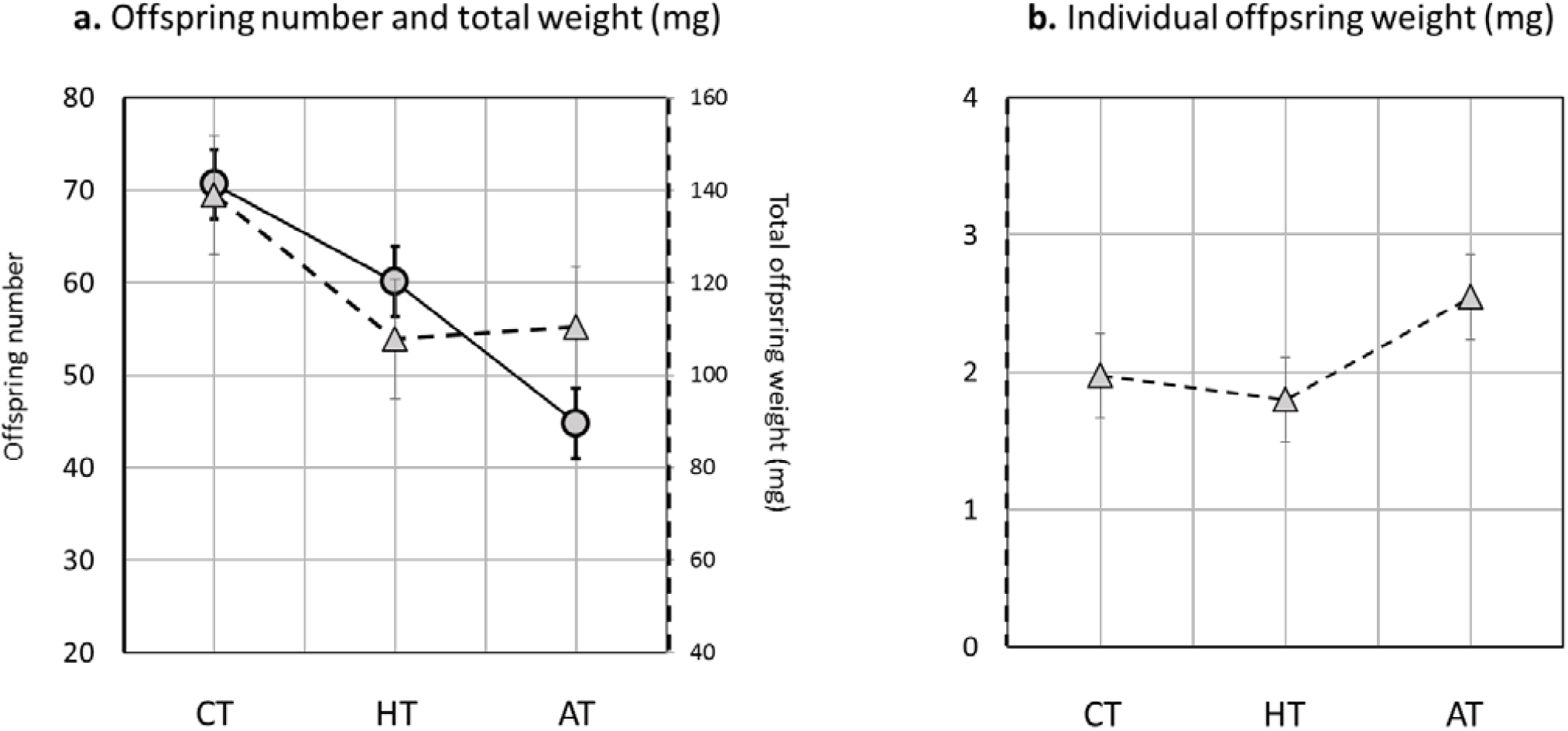
Mean fitness, total offspring weight and individual offspring weight for each treatment. Mean fitness (adult offspring number, circles), mean total offspring weight (triangles, a.), mean individual offspring weight (triangles, b.) and 95% confidence intervals (linear mixed model estimates) are given for the each treatment: Control (CT), Heterogeneous treatment (HT) and Ageing treatment (AT).

**Table 2.** Anova table for a linear mixed model with total offspring weight as a response variable. Type III test. Date of the fitness assay was estimated as random effect.

| Effect | Chi square | df | p.value |
| --- | --- | --- | --- |
| Intercept | 188 | 1 | <0.001 |
| Treatment | 14.1 | 2 | <0.001 |
| Sex | 2.49 | 1 | 0.11 |
| Treatment by Sex | 3.02 | 2 | 0.22 |

**Table 3.** Anova table for a linear mixed model with individual offspring weight as a response variable. Type III test. Date of the fitness assay was estimated as random effect.

| Effect | Chi square | df | p.value |
| --- | --- | --- | --- |
| Intercept | 260 | 1 | <0.001 |
| Treatment | 96.2 | 2 | <0.001 |
| Sex | 3.89 | 1 | 0.049 |
| Treatment by Sex | 3.12 | 2 | 0.21 |

### Sex-specific variance in fitness

Variance was calculated from mean standardized fitness. It is therefore the variance in relative fitness, which represents the opportunity for selection. Variance in relative fitness was larger in males than in females in all three treatments (Figure 3 a and b). The male-bias was largest in the HT, while the CT and AT did not differ from each other (HT-CT:p=0.039, HT-AT: p=0.039, AT-CT=0.45, p-values were obtained from Bayesian posterior distributions, correction for multiple testing was done using the Bonferroni method).

**Figure 3.**
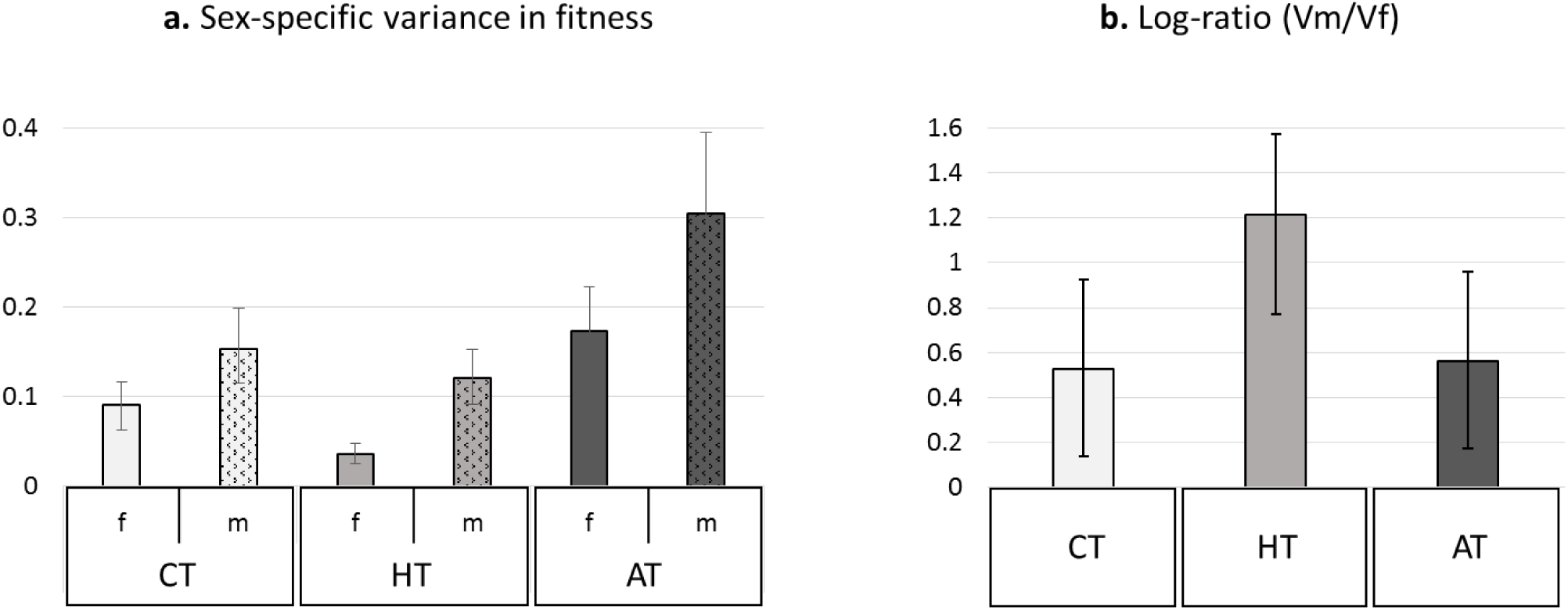
Sex-specific variance in fitness and log-ratio of male over female variance for each treatment. Estimates and 95% confidence interval (Bayesian model estimates) are given for (a) sex specific variance in fitness for females (f, empty bars) and males (m, patterned bars) and for (b) log-ratio of male over female variance log(Vm/Vf). A log-ratio higher than zero indicates male-bias. In (a) and (b), shading refers to the experimental treatment and indicates the level of stress as measured by reduction in mean fitness: clear for CT (no stress), medium shading for HT (intermediate stress) and dark for AT (high stress).

## Discussion

In sexually reproducing species, selection is often measured to be stronger on males that on females, and this sex-bias has often been ascribed to sexual selection acting more on males [21, 22]. This general sex-bias can play an important role in evolution by shaping sexually reproducing populations in many ways, from genetic architecture to mutation load and speed of adaptation. Yet, it is not clear how robust this pattern is to variation in ecological conditions; because the sexes are limited by different resources, variation in sex-specific limiting resources should alter the sex-bias in selection. Here, we used the model species *C. maculatus* to test the hypothesis that limiting female-specific resources should cause a shift towards more female-biased opportunity for selection. However, after challenging females by limiting high-quality oviposition sites (HT) or by delaying age at oviposition (AT), we found that selection remained male-biased and in one case (HT) was even more male-biased than in the control treatment (CT). This result suggests that the trend of male-biased opportunity for selection is robust to variation at least regarding environmental variables studied here. One possible explanation that we discuss below is that selection on males is partly mediated by female choice and therefore reflects selection acting on females as well. Additionally, variance in fitness may not consistently increase in response to stress, which further complicates predictions of how sex-specific selection should behave under stress.

The two experimental treatments, HT and AT, were designed to be challenging and this is confirmed by our results that show how these stressors decrease the mean fitness (adult offspring count) compared to the CT. A general expectation is that variance in fitness should increase under such stressful conditions, as the population is pushed away from its fitness peak [36] and differences among individuals are revealed or magnified [37]. However, as outlined by Hoffmann and Merilä [38], there are scenarios such as severe resource limitation that prevents individuals from expressing their full potential, which allows for a reduction instead of an increase in the opportunity for selection under stress, a prediction that has found some empirical support (reviewed in [37]). This is what we also find here: both male and female opportunity for selection decreased under the HT compared to CT, and female variance decreased proportionally more than male variance resulting in a more male-biased opportunity for selection in that treatment. It is possible that limiting good-quality larval environment in the HT prevented individuals from achieving their full reproductive potential, thereby decreasing variance in relative fitness at the population level, as predicted by Hoffman and Merilä [38]. However, if environmental conditions had imposed a ceiling on reproductive performance, we would have expected to see this reflected in the fitness distributions that should have been more negatively skewed in the HT treatment. We did not observe this (skewness score: CT= −0.38, HT= −0.09, AT= 0,17). In fact, the HT treatment of heterogeneous bean quality should not represent an unsurmountable challenge for female *C. maculatus*, as they are known to be capable of complex oviposition decisions (e.g. [29]).

Alternatively, it is also plausible that, while the HT provided poorer resources that challenged female oviposition strategy and ultimately lowered mean fitness, it may also have removed some of the constraints presented to females in the CT. *C. maculatus* is known for pervasive interlocus sexual conflict, where male mating behavior can substantially lower female lifespan and reproductive success [39]; it is possible that the beans filled with cavities (constituting the majority of the substrate in the HT) offered more hiding opportunities for females to avoid male mating attempts, than fresh beans, as adults easily fit in the bean holes made by previous generations (*personal observation*). There is previous evidence suggesting that more complex laboratory environments could reduce the impact of sexual conflict in *Drosophila melanogaster* [40]. If that is the case here, the HT may have presented females with oviposition challenges but removed or alleviated selection pressure from interlocus sexual conflict. In turn, if the HT made it more difficult for males to find mating partners, this could also explain the stronger male-bias in opportunity for selection in that treatment.

In the AT, the opportunity for selection on females increased, as we expected when imposing a challenge on female oviposition strategy (here, age-at-reproduction). However it also increased proportionally in males, which resulted in a sex-bias similar to the one measured in the CT. We consider several alternative explanations for this result.

Males and females were interacting throughout their lifetime in all of the three treatments, however in the AT, the oviposition was only possible after 48h imposing a constraint particularly to the female reproduction. In a related seed beetle species (*Acanthoscelides obtectus*), experimental work has shown how a selection for a delayed oviposition has resulted in sex-specific evolution of number of life-history traits, including a female-biased elongation of lifespan [41]. However, a constraint to female egg laying is clearly an important factor for males too: there is evidence in *C. maculatus* for last male sperm-precedence [31], which could have favored males in better condition after 48h. This environment could have therefore presented an ageing challenge to both sexes. However, even in that case the different reproductive functions are under selection in the sexes, and the effects of ageing are still expected to be sex-specific with females being more sensitive than males [27, 28].

Another possibility is that the challenge imposed on females by the AT was reverberated onto males through mate choice mechanisms if females confronted to a stressful environment became choosier. The impact of female condition on mate choice has been studied in many systems, however the observations mainly support a weaker mate choice for females in poor conditions (reviewed in [42], and supported by more recent empirical studies [43, 44]). Similarly, in the *A. obtectus* seed beetles mate choice becomes relaxed in females when tested in stressful conditions [45]. These studies indicate that female-specific stress reduces rather than increases the strength of selection imposed on males by female choice. However, a different response could be expected if males can contribute to improve female condition through direct benefits such as nuptial gifts or parental care. In *C. maculatus*, male ejaculate represents a large amount of water, carbohydrates, proteins and peptides, and is sometimes considered a nuptial gift [46, 47] in this aphagous species. It is possible that ageing females would rely more on nutrition and hydration from the contributions of male ejaculate to sustain their reproductive capacity. By imposing selection on delayed reproductive ageing, the AT treatment could have resulted in more stringent mate choice imposed on males that could in turn explain the proportional increase of both the male and female variance in fitness. This mechanism could help to explain the maintenance of male-biased selection even under the limitation of female-specific resources at least in species where mating provides direct resource benefits to females.

### Conclusions

We have shown that there are sex-specific changes in the opportunity for selection in response to different ecological challenges. Although this has been tested before (e.g. [48, 49]), in the present study we placed particular focus on female-specific resource limitation, with the prediction that it would lead to a more female-biased opportunity for selection. This prediction relied on the assumption that resource limitation would generally increase opportunity for selection, which has not been the case for all treatments. Despite the variety of ways in which sex-specific selection responded to our different treatments, selection remained male-biased in all cases, which suggests that this pattern is in fact relatively robust. Moreover, our results from the HT showed that a male-bias in the opportunity for selection can also be driven by a response of females to changes in environmental conditions, which challenges the view that male-bias in selection is generally driven by intense sexual competition in males. While it is not surprising that manipulating variance in fitness of one sex should trigger a response in the other because of the many levels of interactions involved in sexual reproduction, it is rather striking that males remained the more variable sex regardless of the degree of stress on females.

